# Roles of the second-shell amino acid R266 in other members of the MLE subgroup of the enolase superfamily

**DOI:** 10.1101/2022.03.03.481873

**Authors:** Dat P. Truong, Susan Fults, Cristian Davila, Jamison Huddleston, Dakota Brock, Mingzhao Zhu, Jean-Phillipe Pellois, Kenneth G. Hull, Daniel Romo, Frank M. Raushel, Margaret E. Glasner

## Abstract

Catalytic promiscuity is the coincidental ability to catalyze non-biological reactions in the same active site as the native biological reaction. Several lines of evidence show that catalytic promiscuity plays a role in the evolution of new enzyme functions. Thus, studying catalytic promiscuity can help identify structural features that predispose an enzyme to evolve new functions. This study identifies such a pre-adaptive residue in an *N*-succinylamino acid racemase/*o*-succinylbenzoate synthase (NSAR/OSBS) enzymes from the NSAR/OSBS subfamily. Previously, we identified a point mutation, R266Q, in the catalytically promiscuous *Amycolatopsis* sp. T-1-60 NSAR/OSBS that has a deleterious effect on NSAR activity with a lesser effect on OSBS activity (Truong *et al*., in preparation). We demonstrated that R266 was a pre-adaptive feature that enabled the emergence and evolution of NSAR activity in AmyNSAR/OSBS. We examined the role of the residue R266 in the evolution of NSAR activity by examining the effects of the single substitution R266Q in other members of the NSAR/OSBS subfamily including *Enterococcus faecalis* NSAR/OSBS, *Roseiflexus castenholzii* NSAR/OSBS, *Lysinibacillus varians* NSAR/OSBS, and *Listeria innocua* NSAR/OSBS, which have been previously characterized to carry out both OSBS and NSAR activities efficiently. RcNSAR/OSBS, LvNSAR/OSBS, EfNSAR/OSBS, and LiNSAR/OSBS are 49, 48, 32, and 28% identical, respectively, to AmyNSAR/OSBS. We found that while the R266Q mutation decreases NSAR activity more than OSBS activity, as expected, in most NSAR/OSBS members, the differential effects of the R266Q substitution on NSAR and OSBS activities are not as striking as observed in AmyNSAR/OSBS. In some homologs, the R266Q mutation has very deleterious effects on both OSBS and NSAR activities. Furthermore, the mutation unexpectedly decreases OSBS activity more than NSAR activity in LiNSAR/OSBS. Thus, the effects of R266Q on NSAR and OSBS activities depend on differences in sequence context between members of the NSAR/OSBS subfamily, demonstrating the complex role of epistasis in the evolution of NSAR activity in the NSAR/OSBS subfamily.

## Introduction

The evolution of new enzyme functions may require the accumulation of several adaptive mutations. Functional mutations can be found in the first shell amino acids which directly interact with the substrate or second shell amino acids, which interact with the first shell amino acids and so on. While mutations in the first shell amino acids in the active site can be directly responsible for the evolution of enzyme ligand interactions, remote mutations beyond the first shell, including second or third shell and other non-active site residues can contribute to functional adaptation toward new enzymatic functions [1, 2]. Identification of these remote mutations could help us understand the fundamental question in structure-function relationships, which is how do residues throughout the structure interact with each other? Such identifications could also help us understand how new enzymatic activities evolve. However, epistasis limits the ability to correctly identify such mutations. Epistasis occurs when the same mutation has different phenotypic effects in different genetic backgrounds, resulting in the non-additive effects of combinations of mutations [3, 4]. Furthermore, while catalytic residues (or first-shell residues) are required for specific interactions with the substrates and unlikely to exhibit epistasis, non-active site residues (i.e., second- and third-shell residues) are more likely to be prone to epistasis because they could make multiple interactions within different sets of coevolving residues due to the intertwined nature of amino acid network within the enzyme [2].

Previously, we identified a point mutation, R266Q, in the catalytically promiscuous *Amycolatopsis* sp. T-1-60 *N*-succinylamino acid racemase/*o*-succinylbenzoate synthase (AmyNSAR/OSBS) that has a deleterious effect on NSAR activity with a lesser effect on OSBS activity (Truong *et al*., in preparation). AmyNSAR/OSBS belongs to the NSAR/OSBS subfamily, in which many members are catalytically promiscuous and can catalyze both OSBS and NSAR reactions efficiently. The R266Q mutation in AmyNSAR/OSBS profoundly reduces NSAR activity, but only moderately reduces OSBS activity. The second-shell amino acid R266 is close to the catalytic acid/base K263, but it does not contact the substrate. While mutating R266 to glutamine had a minor effect on the enzyme’s structure, the mutation severely decreased the rate of proton exchange between the alpha proton of the NSAR substrate (*N*-succinylphenylglycine, NSPG) and the general acid/base catalyst K263. This mutation is less deleterious for the OSBS reaction because K263 forms a cation-π interaction with the OSBS substrate (2-Succinyl-6-hydroxy-2,4-cyclohexadiene-1-carboxylate, SHCHC) and/or the intermediate, rather than acting as a general acid/base catalyst. We previously showed that R266 was a conserved second-shell residue in the NSAR/OSBS subfamily and not present in other non-promiscuous OSBS subfamilies. We demonstrated that R266 was a pre-adaptive feature that enabled the emergence and evolution of NSAR activity in AmyNSAR/OSBS. However, if R266 is truly pre-adaptive, we expect that mutations at this position will have the same phenotypic effects on the NSAR and OSBS activities in other NSAR/OSBS members.

Phylogenetic and genome context analysis of other members of the NSAR/OSBS subfamily indicates that NSAR activity evolved through promiscuous intermediates (Figure 1). For example, the biological function of the NSAR/OSBS enzymes from many species of *Amycolatopsis*, including Amycolatopsis sp. T-1-60, is expected to be NSAR activity because these species do not require OSBS activity to make menaquinone or there is a separate OSBS gene encoded in the menaquinone operons [5]. The *Enterococcus faecalis* NSAR/OSBS (EfNSAR/OSBS) gene is encoded in a menaquinone operon, indicating that OSBS activity is its biological function [6]. The NSAR/OSBS enzymes from *Listeria innocua, Lysinibacillus varians*, and *Roseiflexus castenholzii* are predicted to be bifunctional, because OSBS activity is required for menaquinone synthesis but the NSAR/OSBS gene is not in the menaquinone operon. Instead, the NSAR/OSBS enzymes from *L. varians* and *R. castenholzii* are encoded in operons with genes from the D- to L-amino acid conversion pathway [5]. The D- to L-amino acid conversion pathway, which was first identified in *Geobacillus kaustophilus*, consists of a succinyltransferase (GNAT superfamily), an NSAR/OSBS, and an L-desuccinylase (M20 family) [7].

**Figure 1.**
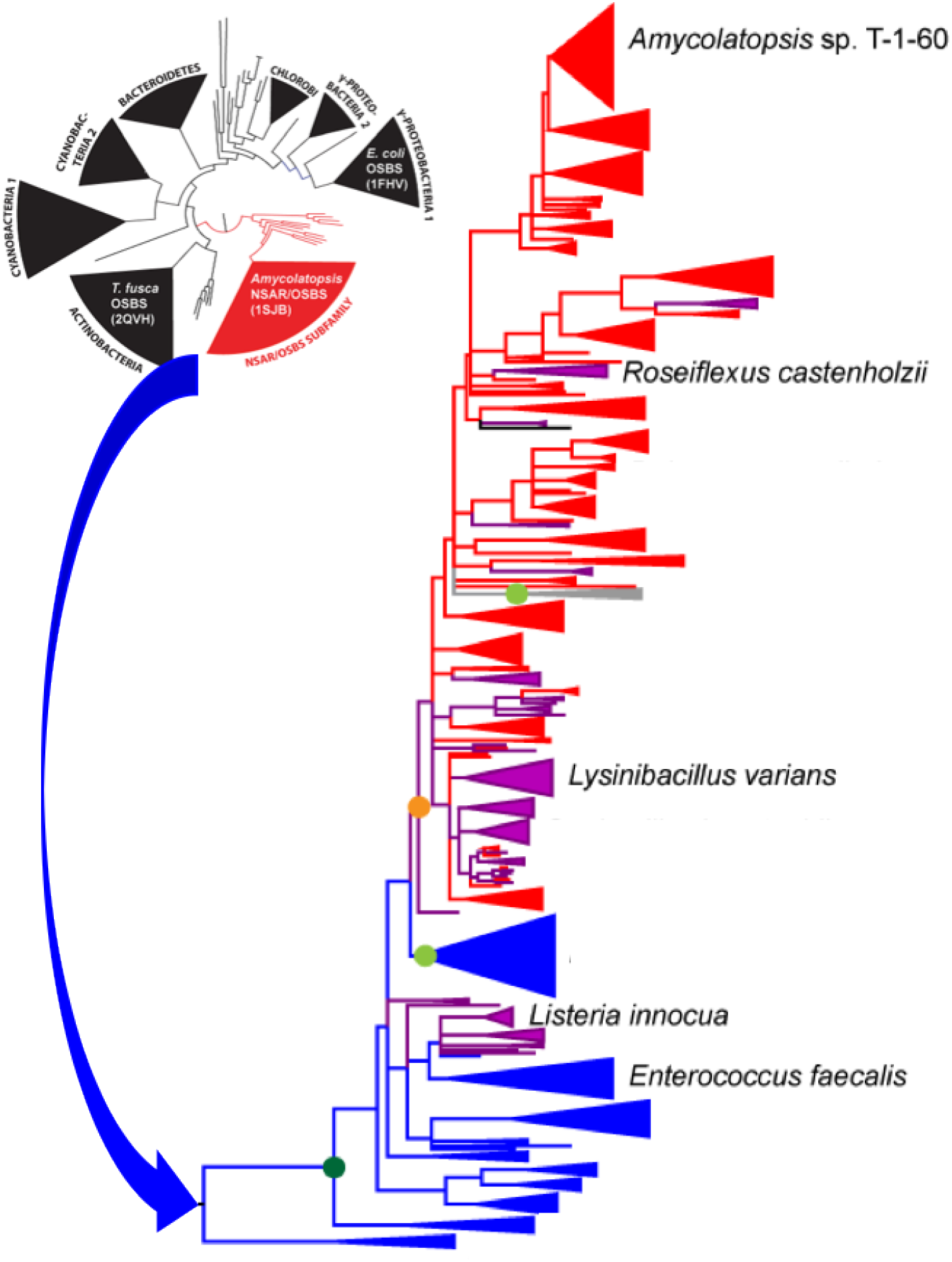
Phylogenetic distribution of the NSAR/OSBS enzymes used in this study. The inset represents several large, divergent OSBS subfamilies, which correspond to the phylum from which the OSBS originated [11]. The blue arrow represents the zoomed-in phylogenetic tree of the NSAR/OSBS subfamily [5]. Blue branches indicate proteins that are encoded in menaquinone operons, indicating that OSBS activity is their biological function. Red branches indicate proteins whose biological function is expected to be NSAR activity because their species do not require OSBS activity to make menaquinone or there is a separate OSBS gene encoded in the menaquinone operon. Purple branches indicate proteins that are predicted to be bifunctional, because OSBS activity is required for menaquinone synthesis but the NSAR/OSBS subfamily gene is not in the menaquinone operon. Many of these proteins are encoded in operons with genes from the D-amino acid conversion pathway.

**Figure 2.**
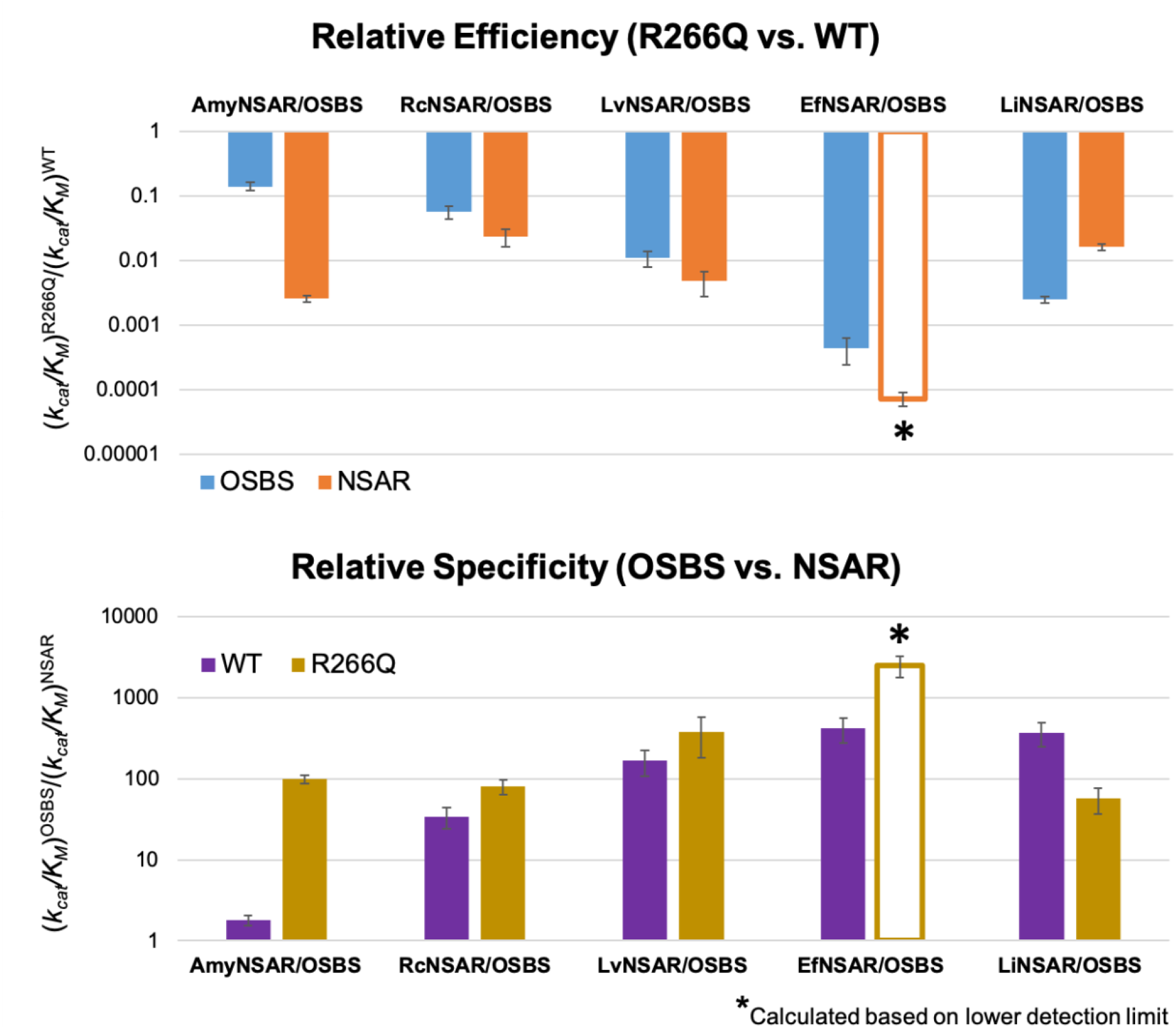
The effects of R266Q in several members of the NSAR/OSBS subfamily. (A) Relative efficiency ratios of R266Q versus WT variants for OSBS (cyan) and NSAR activities (orange). (B). Relative specificity ratio of OSBS activity versus NSAR activity for WT (purple) and R266Q variants (gold). The asterisk indicates that the NSAR activity of this variant was below the detection limit, so *K*_M_^NSAR^ value for EfNSAR/OSBS R266Q was estimated assuming that *K*_M_^NSAR^ is the same in the mutant and wild type, as observed for OSBS activity, and *k_cat_* was estimated as the lower limit of detection.

Sequence divergence of homologous enzymes has an important effect on evolvability of promiscuous and native activities [8]. For example, a study about eight homologs of the L-gamma-glutamyl phosphate (GP) reductase (ProA), which promiscuously catalyzes the N-acetyl-L-glutamyl phosphate (NAGP) reduction, showed that the degree of improvement of the promiscuous NAGP reductase activity achieved by the single mutation E383A varied dramatically and did not correlate with the starting level of the promiscuous activity [8]. The authors speculated that the E383A mutation might remove some steric conflicts so the enzyme could accommodate the binding of NAGP, which is slightly larger than GP [9]. This single mutation in ProA also has differential effects on the native activity of the eight homologs. This study illustrates how the effect of a mutation depends on the sequence and structural contexts in which the mutation occurs (that is, epistatic constraints). Unlike ProA, which exhibits substrate promiscuity, NSAR/OSBS enzymes are catalytically promiscuous. Furthermore, while the mechanism of E383’s effect on substrate specificity of ProA was speculative, the effect of R266 on specificity determination in AmyNSAR/OSBS enzymes has been demonstrated (Truong *et al*., in preparation). Members of the NSAR/OSBS subfamily are moderately divergent, generally sharing >40% sequence identity, while having variation in their relative OSBS and NSAR activities and differences in their biological function [5]. This raises the question that R266 might not have the same role in different sequence backgrounds.

Here, we examine the role of the residue R266 in the evolution of NSAR activity by examining the effects of the single substitution R266Q in other members of the NSAR/OSBS subfamily. We made an arginine-to-glutamine substitution at the homologous position of EfNSAR/OSBS, *Roseiflexus castenholzii* NSAR/OSBS (RcNSAR/OSBS), *Lysinibacillus varians* NSAR/OSBS (LvNSAR/OSBS), and *Listeria innocua* NSAR/OSBS (LiNSAR/OSBS), which have been previously characterized. These enzymes efficiently carry out both OSBS and NSAR activities [5, 10]. RcNSAR/OSBS, LvNSAR/OSBS, EfNSAR/OSBS, and LiNSAR/OSBS are 49, 48, 32, and 28% identical, respectively, to AmyNSAR/OSBS. We found that while the R266Q mutation decreases NSAR activity more than OSBS activity, as expected, in most NSAR/OSBS members, the differential effects of the R266Q substitution on NSAR and OSBS activities are not as striking as observed in AmyNSAR/OSBS. In some homologs, the R266Q mutation has very deleterious effects on both OSBS and NSAR activities. Furthermore, the mutation unexpectedly decreases OSBS activity more than NSAR activity in LiNSAR/OSBS. Thus, the effects of R266Q on NSAR and OSBS activities depend on differences in sequence context between members of the NSAR/OSBS subfamily, demonstrating the complex role of epistasis in the evolution of NSAR activity in the NSAR/OSBS subfamily.

## Materials and Methods

### Mutagenesis

Site-directed mutagenesis was performed by Q5 mutagenesis (New England BioLabs). The templates for mutagenesis included: the gene encoding *Enterococcus faecalis* NSAR/OSBS (EfNSAR/OSBS) (UniProt entry: Q838J7) and the gene encoding for *Listeria innocua* NSAR/OSBS (LiNSAR/OSBS) (UniProt entry: Q927X3), which were cloned into a pET15b vector (Novagen) (gifts from J.A Gerlt, University of Illinois, Urbana, IL); the gene encoding *Lysinibacillus varians* GY32 NSAR/OSBS (LvNSAR/OSBS) (UniProt entry: X2GR01) and the gene encoding *Roseiflexus castenholzii* HLO8 NSAR/OSBS (RcNSAR/OSBS) (UniProt entry: A7NLX0) which were cloned into a modified pET21a vector pMCSG7 or pMSCSG8, which encodes an N-terminal His6 tag, via ligation-independent cloning [12]. The genes encoding *E. coli* Dipeptide Epimerase (EcDE) (UniProt entry: P51981) and *B. subtilis* Dipeptide Epimerase (BsDE) (UniProt entry: O34508) were cloned into a modified pET15b vector, which encodes an N-terminal His10 tag (gifts from J.A Gerlt, University of Illinois, Urbana, IL). Mutations were confirmed by sequencing in both directions (Eurofins Genomics LLC). The primers used for mutagenesis were designed using NEBaseChanger, NEB’s online design software (NEBasechanger.com) and are shown in the Table 1.

**Table 1:**
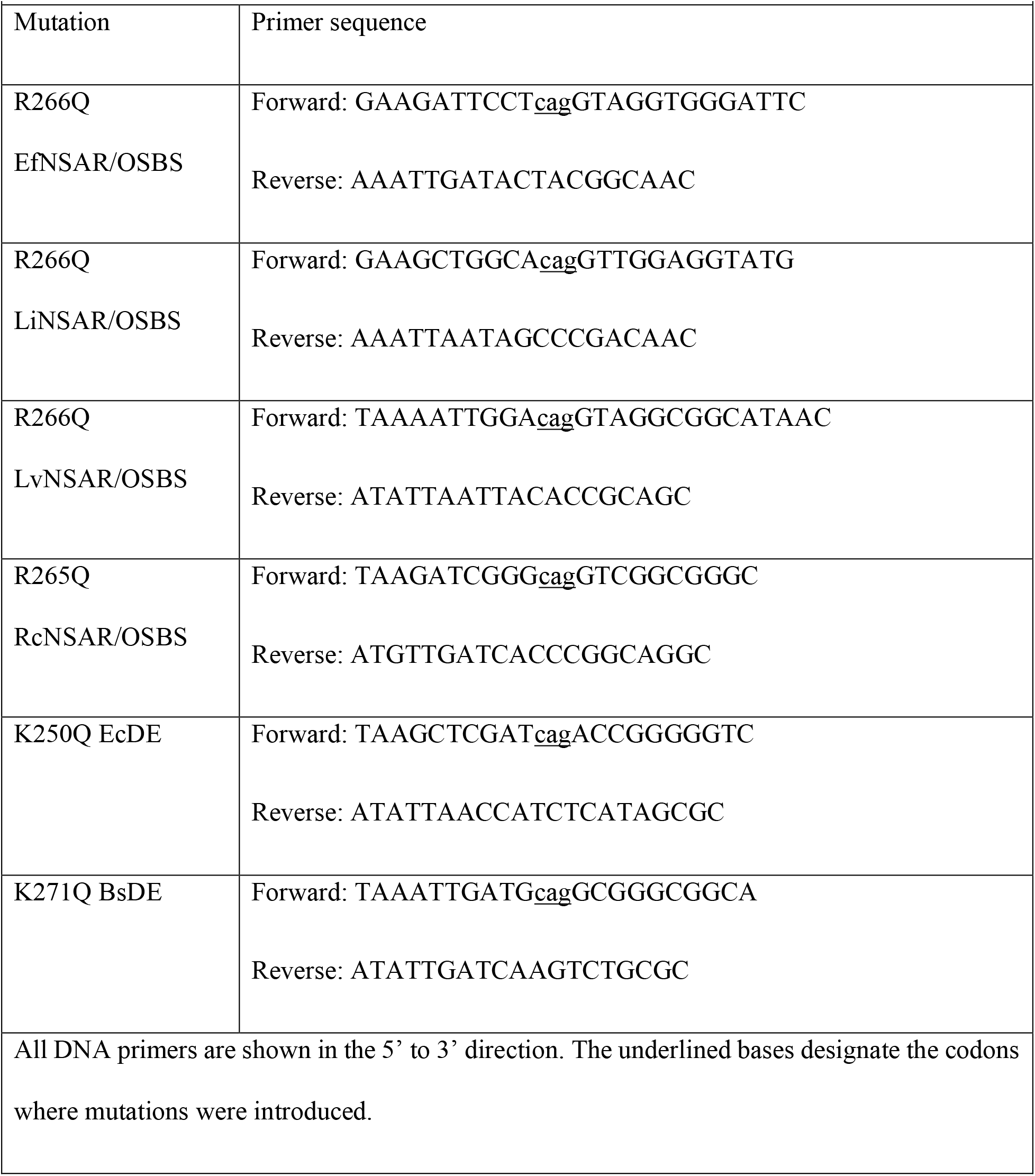
Primers used for mutagenesis of different NSAR/OSBS enzymes

### Protein Production

Proteins were expressed in *E. coli* strain BW 25113 (*menC::kan*, DE3) to ensure that OSBS from the host cell would not contaminate the purified proteins expressed from the plasmid [11]. Cultures were grown overnight at 37°C in LB media containing carbenicillin and kanamycin at a final concentration of 50 μg/mL with no induction, then harvested by centrifugation. The cell pellet was resuspended in 10 mM Tris (pH 8.0), 500 mM NaCl, 5 mM imidazole, 0.4 mM phenylmethylsulfonyl fluoride (PMSF), and 10 μg/mL DNase I. The supernatant was collected after centrifugation and filtered using a 0.22 μm Steriflip filter (Millipore). The protein was loaded into a 5 mL HisTrap FF column charged with Ni^2+^ (GE Healthcare). The protein was eluted using a buffer containing 10 mM tris (pH 8.0), 500 mM NaCl, and 500 mM imidazole with a step to 15% elution buffer to elute loosely bound proteins, followed by a linear gradient to 100% elution buffer over 20 column volumes. Fractions containing the proteins were identified by SDS-PAGE and concentrated using a Vivaspin Turbo 15 centrifuge filter with a 10 kDa molecular weight cutoff (Sartorius). Glycerol was added to a final concentration of 25%, and the purified proteins were stored at −20 °C.

### OSBS Assay

2-Succinyl-6-hydroxy-2,4-cyclohexadiene-1-carboxylate (SHCHC) was synthesized from chorismate and α-ketoglutarate as described previously [11]. The enzyme was assayed in 50 mM Tris (pH 8.0) and 0.1 mM MnCl_2_ with various SHCHC concentrations. The reactions were monitored by a SpectraMax Plus384 plate reader (Molecular Devices) at 310 nm an/d at 25 °C. The disappearance of SHCHC (Δ_ε_ = −2400 M^−1^ cm^−1^) was measured as a function of time [13, 14]. The initial rates were determined by fitting the linear portion of the data in Microsoft Excel, and the initial rates at different substrate concentrations were fit to the Michaelis-Menten equation using Prism (GraphPad).

### NSAR Assay

L- and D-*N*-Succinylphenylglycine (L- and D-NSPG) were synthesized as described previously [15]. The enzymes were assayed in 200 mM Tris (pH 8.0) and 0.1 mM MnCl_2_ with various L- or D-*N*-succinylphenylglycine concentrations. The reactions were carried out in a sample cell with a 5 cm path length. The change in the optical rotation of the substrate was monitored by a Jasco P-2000 polarimeter at 405 nm and at 25 °C. Measurements were taken using a 10 s integration time and read every 30 seconds. The specific rotation value at 405 nm of L-NSPG is 6.54 deg M^−1^ cm^−1^ and of D-NSPG is 6.22 deg M^−1^ cm^−1^ [15]. The rates were determined as described above.

### Dipeptide Epimerase Activity

L-Ala-L-Glu was commercially purchased (Chem Impex Int’l, Inc) and L-Ala-D-Glu was synthesized as described previously [16]. The dipeptide epimerase enzyme was assayed in 50 mM Tris (pH 8.0), 10 mM MgCl_2_ with various L-Ala-L-Glu concentrations. The reactions were carried out in a cell with a 5 cm path length in a total volume of 1.4 mL. The change in the optical rotation of the substrate was monitored by a Jasco P-2000 polarimeter at 365 nm and at 25 °C. Measurements were taken using a 10 s integration time and reading every 30 seconds for a total of 30 minutes. The initial rates were determined by fitting the linear portion of the data in Microsoft Excel, and the initial rates at different substrate concentrations were fit to the Michaelis-Menten equation using Prism, GraphPad.

### Isotopic Exchange Experiments Using ^1^H NMR Spectroscopy

The NSAR/OSBS and dipeptide epimerase variants were exchanged into D2O using a Vivaspin Turbo 15 centrifuge filter (Sartorius). A 10 mL aliquot of protein was concentrated to 1 mL; 9 mL of D2O was added, and the protein solution was again concentrated to 1 mL. This process was repeated three times to maximize the exchange. Each NSAR reaction contained 20 mM L- or D-NSPG (pD 8.0), 50 mM Tris (pD 8.0), 0.1 mM MnCl_2_, and NSAR/OSBS variants in D2O. The intensity of the α-proton was monitored as it is exchanged with deuterium over time by ^1^H NMR (500 MHz Bruker NMR spectrometer). The peak for the α-proton (δ = 5.15 ppm) was integrated relative to that of the five aromatic protons (δ = 7.40 ppm). The relative peak area was converted to concentration based on the initial substrate concentration. The slopes of the plots of the NSPG substrate concentration as a function of time were fit to a line to obtain the isotopic exchange rates (*k_ex_*).

Each dipeptide epimerase reaction contained 10 mM L-Ala-L-Glu or L-Ala-D-Glu (pD 8.0), 20 mM Tris (pD 8.0), 0.1 mM McCl_2_, and dipeptide epimerase enzymes in D2O. The intensity of the α-proton was monitored as it is exchanged with deuterium over time by ^1^H NMR (500 MHz Bruker NMR spectrometer). The peak for the α-proton (δ = 4.08 ppm) was integrated relative to that of the H_α_ of Ala (δ = 3.95 ppm). The relative peak area was converted to concentration based on the initial substrate concentration. The slopes of the plots the L-Ala-L/D-Glu substrate concentration as a function of time were fit to a line to obtain the isotopic exchange rates (*k_ex_*).

## Results

### The roles of R266 in other members of the NSAR/OSBS subfamily

Previously, we demonstrated that R266 was a pre-adaptive feature that enabled the emergence and evolution of NSAR activity in AmyNSAR/OSBS (Truong *et al*., in preparation). However, if R266 is truly pre-adaptive, we expect that the R266Q mutation will have the same phenotypic effects on the NSAR and OSBS activities in other NSAR/OSBS members. To determine if R266 has the same effect on the activities in other members of the NSAR/OSBS subfamily, we made an arginine-to-glutamine substitution at the homologous position 266 in *Enterococcus faecalis* NSAR/OSBS (EfNSAR/OSBS), *Roseiflexus castenholzii* NSAR/OSBS (RcNSAR/OSBS), *Lysinibacillus varians* NSAR/OSBS (LvNSAR/OSBS), and *Listeria innocua* NSAR/OSBS (LiNSAR/OSBS). These enzymes efficiently carry out both OSBS and NSAR activities, with a much stronger preference toward OSBS activity (Table 2). We expected that the R266Q mutation would decrease NSAR activity more than OSBS activity, as observed in AmyNSAR/OSBS. Overall, the R266Q mutation decreases NSAR activity more than OSBS activity in all variants except LiNSAR/OSBS. However, the R266Q mutation in those enzymes does not affect the relative specificity as dramatically as in AmyNSAR/OSBS. In AmyNSAR/OSBS, while the R266Q mutation has no effect on either K_M_^OSBS^ and K_M_^NSAR^, it only decreased *k*_cat_^OSBS^ by 7-fold and significantly decreased *k*_cat_^NSAR^ by ~470-fold (Truong *et al*., in preparation). In contrast, the R266Q mutation in other NSAR/OSBS enzymes decreased *k_cat_*^OSBS^ by ~7-379-fold and decreased *k_cat_*^NSAR^ ranged from ~10-fold to undetectable, while generally having lesser or no effect on either K_M_^OSBS^ and K_M_^NSAR^.

**Table 2:**
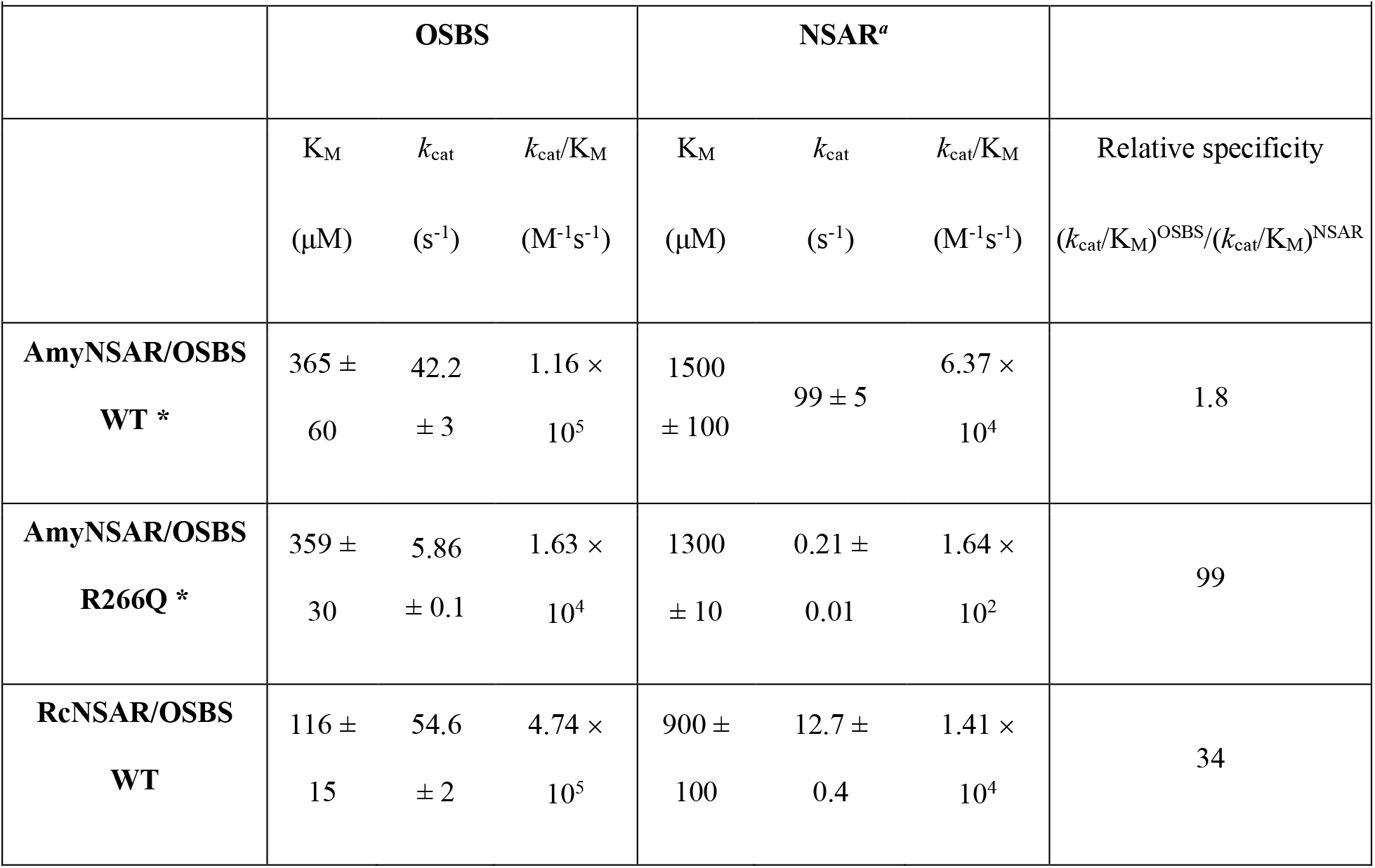
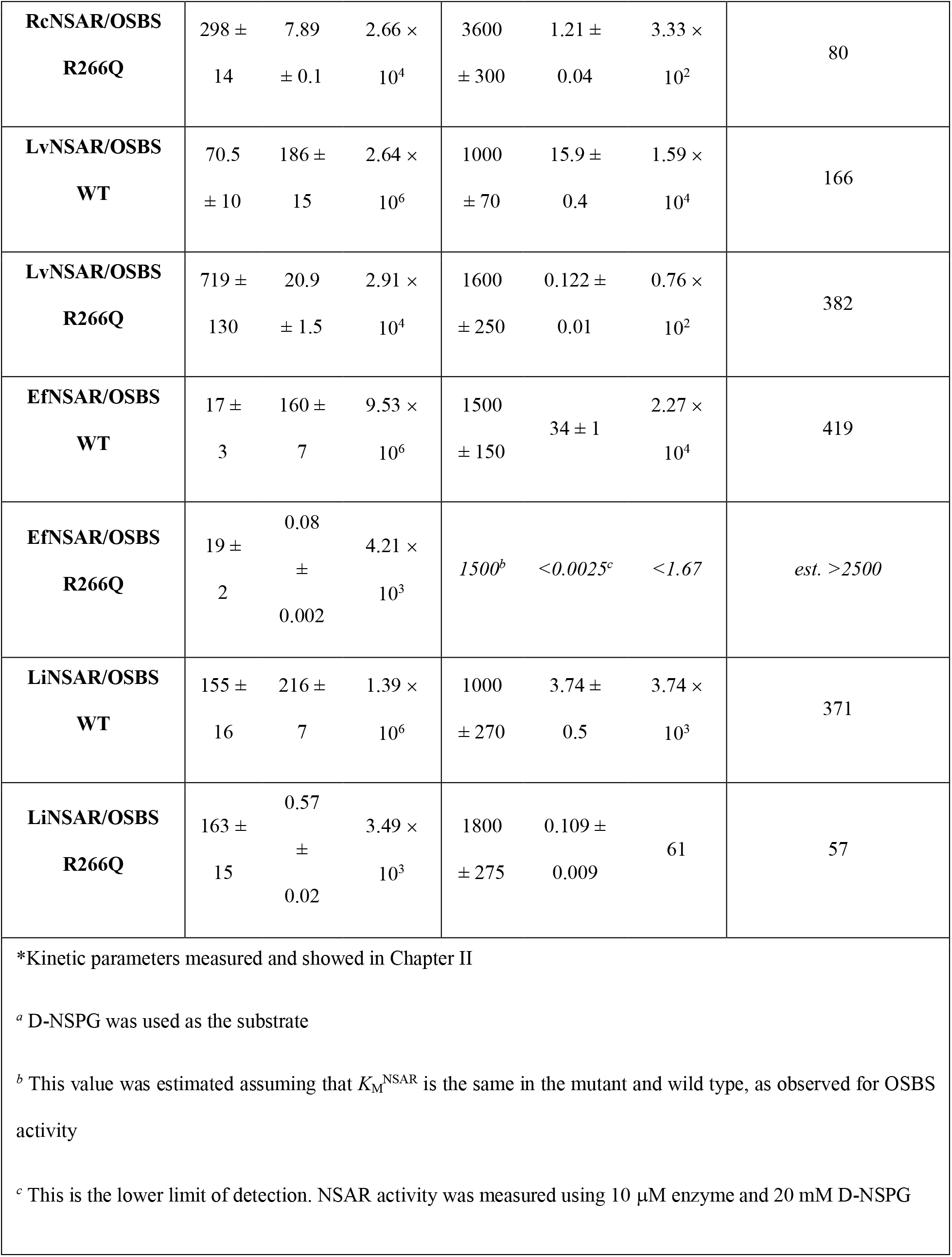
Kinetic constants for WT and R266Q variants of members of the NSAR/OSBS subfamily

|  |  |  |  |  |  |  |  |
| --- | --- | --- | --- | --- | --- | --- | --- |
|  | **OSBS** |  |  | **NSAR<sup>a</sup>** |  |  |  |
| | $K_M$ | $k_{cat}$ | $k_{cat}/K_M$ | $K_M$ | $k_{cat}$ | $k_{cat}/K_M$ | Relative specificity |
| | ($\mu M$) | ($s^{-1}$) | ($M^{-1}s^{-1}$) | ($\mu M$) | ($s^{-1}$) | ($M^{-1}s^{-1}$) | $(k_{cat}/K_M)^{OSBS}/(k_{cat}/K_M)^{NSAR}$ |
| **AmyNSAR/OSBS**<br>**WT \*** | $365 \pm 60$ | $42.2 \pm 3$ | $1.16 \times 10^5$ | $1500 \pm 100$ | $99 \pm 5$ | $6.37 \times 10^4$ | 1.8 |
| **AmyNSAR/OSBS**<br>**R266Q \*** | $359 \pm 30$ | $5.86 \pm 0.1$ | $1.63 \times 10^4$ | $1300 \pm 10$ | $0.21 \pm 0.01$ | $1.64 \times 10^2$ | 99 |
| **RcNSAR/OSBS**<br>**WT** | $116 \pm 15$ | $54.6 \pm 2$ | $4.74 \times 10^5$ | $900 \pm 100$ | $12.7 \pm 0.4$ | $1.41 \times 10^4$ | 34 |
| **RcNSAR/OSBS**<br>**R266Q** | 298 ±<br>14 | 7.89<br>± 0.1 | 2.66 ×<br>10<sup>4</sup> | 3600<br>± 300 | 1.21 ±<br>0.04 | 3.33 ×<br>10<sup>2</sup> | 80 |
| **LvNSAR/OSBS**<br>**WT** | 70.5<br>± 10 | 186 ±<br>15 | 2.64 ×<br>10<sup>6</sup> | 1000<br>± 70 | 15.9 ±<br>0.4 | 1.59 ×<br>10<sup>4</sup> | 166 |
| **LvNSAR/OSBS**<br>**R266Q** | 719 ±<br>130 | 20.9<br>± 1.5 | 2.91 ×<br>10<sup>4</sup> | 1600<br>± 250 | 0.122 ±<br>0.01 | 0.76 ×<br>10<sup>2</sup> | 382 |
| **EfNSAR/OSBS**<br>**WT** | 17 ±<br>3 | 160 ±<br>7 | 9.53 ×<br>10<sup>6</sup> | 1500<br>± 150 | 34 ± 1 | 2.27 ×<br>10<sup>4</sup> | 419 |
| **EfNSAR/OSBS**<br>**R266Q** | 19 ±<br>2 | 0.08<br>±<br>0.002 | 4.21 ×<br>10<sup>3</sup> | 1500<sup>b</sup> | <0.0025<sup>c</sup> | <1.67 | *est. >2500* |
| **LiNSAR/OSBS**<br>**WT** | 155 ±<br>16 | 216 ±<br>7 | 1.39 ×<br>10<sup>6</sup> | 1000<br>± 270 | 3.74 ±<br>0.5 | 3.74 ×<br>10<sup>3</sup> | 371 |
| **LiNSAR/OSBS**<br>**R266Q** | 163 ±<br>15 | 0.57<br>±<br>0.02 | 3.49 ×<br>10<sup>3</sup> | 1800<br>± 275 | 0.109 ±<br>0.009 | 61 | 57 |

\*Kinetic parameters measured and showed in Chapter II
<sup>a</sup> D-NSPG was used as the substrate
<sup>b</sup> This value was estimated assuming that $K_M^{NSAR}$ is the same in the mutant and wild type, as observed for OSBS activity
<sup>c</sup> This is the lower limit of detection. NSAR activity was measured using 10 µM enzyme and 20 mM D-NSPG

To determine if the R266Q mutation decreases the reactivity of the catalytic K263 as observed in AmyNSAR/OSBS R266Q, we used ^1^H NMR spectroscopy to measure the exchange rate (*k_ex_*) between the alpha proton of D- or L-NSPG and the catalytic lysines (Table 3). This experiment allows us to observe the rate of the first step of the NSAR reaction. Previously, we showed that the R266Q mutation slightly decreased the *k_ex_* value of K163, while it remarkably decreased the *k_ex_* value of K263 by 1340-fold in AmyNSAR/OSBS (Truong *et al*., in preparation). We expected that the R266Q mutation would decrease the *k_ex_* value of K263 with a lesser effect on the *k_ex_* value of K163 in other NSAR/OSBS enzymes, as observed in AmyNSAR/OSBS. Overall, the R266Q mutation decreased the *k_ex_* value of K263 more than that of K163 in all homologs, as expected (Table 2). However, in several cases, the mutation also significantly decreased the *k_ex_* values of K163 (EfNSAR/OSBS, LvNSAR/OSBS, and LiNSAR/OSBS), contrasting with our expectation. We will discuss the effects of the R266Q mutation on OSBS and NSAR activities and the reactivity of the catalytic lysines in each individual enzyme below.

**Table 3:** Deuterium-hydrogen exchange rate (*k_ex_*) of other NSAR/OSBS variants

|  |  |  |  |
| --- | --- | --- | --- |
|  | **AmyNSAR/OSBS WT** | **AmyNSAR/OSBS R266Q** | **Fold change** |
| D-NSPG and K163 | 90 s<sup>-1</sup> | 25 s<sup>-1</sup> | 3.6 |
| L-NSPG and K263 | 134 s<sup>-1</sup> | 0.1 s<sup>-1</sup> | 1340 |
|  | **RcNSAR/OSBS WT** | **RcNSAR/OSBS R266Q** | **Fold change** |
| D-NSPG and K163 | 11 s<sup>-1</sup> | 10 s<sup>-1</sup> | 1.1 |
| L-NSPG and K263 | 22 s<sup>-1</sup> | 0.65 s<sup>-1</sup> | 34 |
|  | **LvNSAR/OSBS WT** | **LvNSAR/OSBS R266Q** | **Fold change** |
| D-NSPG and K163 | 156 s<sup>-1</sup> | 2 s<sup>-1</sup> | 78 |
| L-NSPG and K263 | 43 s<sup>-1</sup> | 0.4 s<sup>-1</sup> | 108 |
|  | **LiNSAR/OSBS WT** | **LiNSAR/OSBS R266Q** | **Fold change** |
| D-NSPG and K163 | 68 s<sup>-1</sup> | 5 s<sup>-1</sup> | 14 |
| L-NSPG and K263 | 6 s<sup>-1</sup> | 0.2 s<sup>-1</sup> | 30 |
|  | **EfNSAR/OSBS WT** | **EfNSAR/OSBS R266Q** | **Fold change** |
| D-NSPG and K163 | 50 s<sup>-1</sup> | 0.02 s<sup>-1</sup> | 2500 |
| L-NSPG and K263 | 2.7 s<sup>-1</sup> | <0.002 s<sup>-1</sup> <sup>a</sup> | >1350 |
| <sup>a</sup> This value is the lower detection limit. |  |  |  |

RcNSAR/OSBS has a lower relative specificity than the other enzymes and is the most similar to AmyNSAR/OSBS in that respect. Likewise, the effect of the R266Q mutation in RcNSAR/OSBS is also the most similar to its effect on AmyNSAR/OSBS. RcNSAR/OSBS R266Q mutant slightly increased *K*_M_^OSBS^ and decreased *k_cat_*^OSBS^ by 7-fold, resulting in a ~17-fold drop in *k_cat_/K*_M_^OSBS^, while it increased *K*_M_^NSAR^ by 4-fold and decreased *k_cat_*^NSAR^ by 10-fold, resulting in a ~43-fold drop in *K_cat_/K*_M_^NSAR^ agreeing with our expectation. In RcNSAR/OSBS, the R266Q mutation did not affect the *k_ex_* value of K163 at all while it decreased the *k_ex_* value of K263 by 34-fold. The effects on the reactivity of K163 and K263 in RcNSAR/OSBS are most similar to the effects observed in AmyNSAR/OSBS. The decrease of the *k_ex_* value of K263 is the same order of magnitude with the drop in the *k_cat_*/*K*_M_^NSAR^ in RcNSAR/OSBS. This agrees with our hypothesis that R266 is a pre-adaptive feature and required to modulate the reactivity of K263, as observed in AmyNSAR/OSBS.

In LvNSAR/OSBS, the R266Q mutation unexpectedly increased *K*_M_^OSBS^ by 10-fold and decreased *k_cat_*^OSBS^ by 9-fold, resulting in a ~90-fold decrease in *k_cat_/K*_M_^OSBS^, while it approximately has no effect on *K*_M_^NSAR^ and decreased *k_cat_*^NSAR^ by 130-fold, resulting in a ~209-fold decrease in k_cat_/*K*_M_^NSAR^. We did not expect the R266Q mutation to affect *K*_M_ values in the NSAR/OSBS enzymes, and it was surprising to see that it significantly increased *K*_M_^OSBS^ in LvNSAR/OSBS. This suggests that the R266Q mutation might cause a large structural effect and distort the active site of LvNSAR/OSBS, affecting the binding of the OSBS substrate. In LvNSAR/OSBS, the R266Q mutation decreased the *k_ex_* values of both K163 and K263 by 78- and 108-fold, respectively, approximately the same order with the decrease in both *k_cat_*/*K*_M_^OSBS^ and *k_cat_*/*K*_M_^NSAR^ in LvNSAR/OSBS R266Q. The R266Q mutation decreased the reactivity of both K163 and K263, suggesting that the mutation also has a large structural effect on the active site.

Of the four additional NSAR/OSBS enzymes, EfNSAR/OSBS has the highest relative specificity, with a much stronger preference toward OSBS activity. In EfNSAR/OSBS, while the R266Q mutation dramatically decreased *k_cat_*^OSBS^ by ~2000-fold without affecting *K*_M_^OSBS^, the EfNSAR/OSBS R266Q mutant had no detectable NSAR activity even with 10 μM enzyme. The *k_cat_/K*_M_^NSAR^ value dropped >10^4^-fold, assuming *K*_M_^NSAR^ was unchanged from the WT enzyme, as observed in the OSBS reaction. Although the R266Q mutation decreased NSAR activity more than OSBS in EfNSAR/OSBS, it had the most deleterious effects on both OSBS and NSAR reactions out of all the enzymes. In EfNSAR/OSBS, the R266Q mutation also had the strongest deleterious effect on the *k_ex_* values. While the *k_ex_* value of K163 decreased dramatically by 2500-fold, the *k_ex_* value of K263 was below the detection limit, which explained why we could not detect any NSAR activity in EfNSAR/OSBS R266Q. Thus, the fold change of the *k_ex_* value of K263 by the R266Q substitution in this enzyme was estimated based on the lower limit of detection, resulting in an uncertainty in the differential effect of the R226Q on the catalytic lysines. However, the drop in the *k_ex_* value of K163 is in the same order of magnitude with the drop of *k_cat_*^OSBS^, explaining the decrease of OSBS activity in EfNSAR/OSBS R266Q.

Finally, in LiNSAR/OSBS, the R266Q mutation significantly decreased *k_cat_*^OSBS^ by ~379-fold with no effect on *K*_M_^OSBS^, while it approximately has no effect on *K*_M_^NSAR^ and decreased *k_cat_*^NSAR^ by only 37-fold, resulting in only a 61-fold decrease in *k_cat_/K*_M_^NSAR^. Unlike other enzymes, LiNSAR/OSBS R266Q unexpectedly decreased OSBS activity more than NSAR activity, contrasting with our prediction. However, it is also worth noting that LiNSAR/OSBS shares the lowest sequence identity with AmyNSAR/OSBS, and LiNSAR/OSBS WT already has a much lower level of NSAR activity compared to a much higher level of OSBS activity. Because LiNSAR/OSBS is much less optimal to carry out NSAR activity than other NSAR/OSBS enzymes, introducing the R266Q mutation did not necessarily have as dramatic an effect on NSAR activity as in other enzymes. Furthermore, in LiNSAR/OSBS, the R266Q mutation decreased the *k_ex_* values of both K163 and K263 by 14- and 30-fold, respectively. Even though the R266Q mutation decreased OSBS activity more than NSAR activity in LiNSAR/OSBS, the reactivity of K263 is slightly decreased more than that of K163. The decrease of the *k_ex_* value of K263 is the same order of magnitude with the drop in the *k_cat_*^NSAR^ in LiNSAR/OSBS. However, the *k_ex_* of K163 data did not explain the drop in *k_cat_*^OSBS^.

### Effects of R266 on stability of NSAR/OSBS variants

Because the protein yield from purifying some NSAR/OSBS mutants was lower than the WT enzymes, we measured the thermostability of the NSAR/OSBS variants using differential scanning fluorimetry (DSF). DSF was previously described as a good method to measure the melting temperature (*T_m_*) of proteins [17]. The NSAR/OSBS enzymes in this study are multimeric proteins, and although we did not observe multiple unfolding transitions with the DSF experiments, the *T_m_* should be considered apparent. Overall, the R266Q mutation has a major destabilizing effect in all NSAR/OSBS enzymes, except RcNSAR/OSBS (Table 4). RcNSAR/OSBS R266Q’s *T_m_* is 6.2 °C higher than the WT enzyme. The *T_m_* values in other NSAR/OSBS R266Q mutations are 7.6 – 18 °C lower than the WT enzymes, suggesting that the R266Q mutation has a major destabilizing effect. LvNSAR/OSBS R266Q has the most severe destabilizing effect (18 °C lower than the WT enzyme), suggesting that this was the primary reason that the mutation decreased the proton exchange rate of both K263 and K163 in LvNSAR/OSBS R266Q. It was previously shown that the R266Q mutation has only a minor effect on protein stability in AmyNSAR/OSBS (Truong *et al*., *in preparation*). These data suggest that R266 also contributes to protein stability in the NSAR/OSBS subfamily.

**Table 4:** Estimates of apparent *T_m_* of NSAR/OSBS variants.

| **Table 4:** Estimates of apparent $T_m$ of NSAR/OSBS variants. | |
| --- | --- |
| **NSAR/OSBS Variants** | **$T_m$ (°C)** |
| RcNSAR/OSBS WT | 47.6 (± 0.2) |
| RcNSAR/OSBS R266Q | 53.8 (± 0.3) |
| LvNSAR/OSBS WT | 60.8 (± 0.6) |
| LvNSAR/OSBS R266Q | 42.8 (± 0.8) |
| LiNSAR/OSBS WT | 54.9 (± 0.4) |
| LiNSAR/OSBS R266Q | 42.9 (± 0.6) |
| EfNSAR/OSBS WT | 55.5 (± 0.1) |
| EfNSAR/OSBS R266Q | 47.9 (± 0.1) |

The observed effects of R266Q on OSBS and NSAR activities in different members of the NSAR/OSBS subfamily are due to the different sequence contexts of each NSAR/OSBS enzyme, or epistasis. We observed some correlation between sequence divergence of the NSAR/OSBS enzymes and the phenotypic effects on NSAR and OSBS by the single point mutation R266Q. RcNSAR/OSBS, LvNSAR/OSBS, EfNSAR/OSBS, and LiNSAR/OSBS are 49, 45, 40, and 37% identical, respectively, to AmyNSAR/OSBS. RcNSAR/OSBS shares the highest sequence identity with AmyNSAR/OSBS, and the effects of the R266Q mutation have the most similar phenotypic effects with AmyNSAR/OSBS R266Q. On the other hand, LiNSAR/OSBS shares the least identity with AmyNSAR/OSBS, and LiNSAR/OSBS R266Q has the opposite phenotypic effect on NSAR and OSBS activities compared with AmyNSAR/OSBS R266Q. Here, we can see that epistasis plays an important role in the evolution of NSAR activity in the NSAR/OSBS subfamily. Due to the complexity of epistasis, the phenotypic effects on activities of the same amino acid substitution and the roles of the conserved residues, such as R266, cannot be always assumed to be the same in different NSAR/OSBS homologs.

We attempted to understand the effects of the R266Q mutation by examining the environment surrounding the R266 residue in the crystal structures of EfNSAR/OSBS and LiNSAR/OSBS and comparing that to the structure of AmyNSAR/OSBS [18]. However, only the apo-structure of EfNSAR/OSBS (PDB ID 1WUE) and Mg^2+^ ion-bound structure of LiNSAR/OSBS (PDB ID 1WUF) are available [10]. Overall, the environments within 5 Å surrounding the R266 residue are very similar (Figure 3). Only two positions differ significantly in the aligned structures. First, position 243 in AmyNSAR/OSBS is a valine, while it is substituted with an arginine in both EfNSAR/OSBS and LiNSAR/OSBS. Second, position 97 in AmyNSAR/OSBS is an arginine while it is a tryptophan in EfNSAR/OSBS and a glutamate in LiNSAR/OSBS. R243 and E97 form a salt bridge in LiNSAR/OSBS while in EfNSAR/OSBS, R243 and W97 might form a cation-π interaction. These interactions electrostatically contribute to the active site, potentially making the area around R266 more rigid, and less able to tolerate the R266Q substitution in these enzymes. These interactions are not present in AmyNSAR/OSBS and could account for the different effects of the R266Q mutation. These observations indicate that small differences surrounding R266 in different NSAR/OSBS homologs might contribute to the different effects of the R266Q mutation. Because of sequence divergence of the NSAR/OSBS enzymes in this study, other residues that are more remote from R266 could also potentially influence the effects of the R266Q mutation. Identifications of these residues that influence R266 remotely are necessary to fully understand the evolution of NSAR activity in the NSAR/OSBS subfamily.

**Figure 3.**
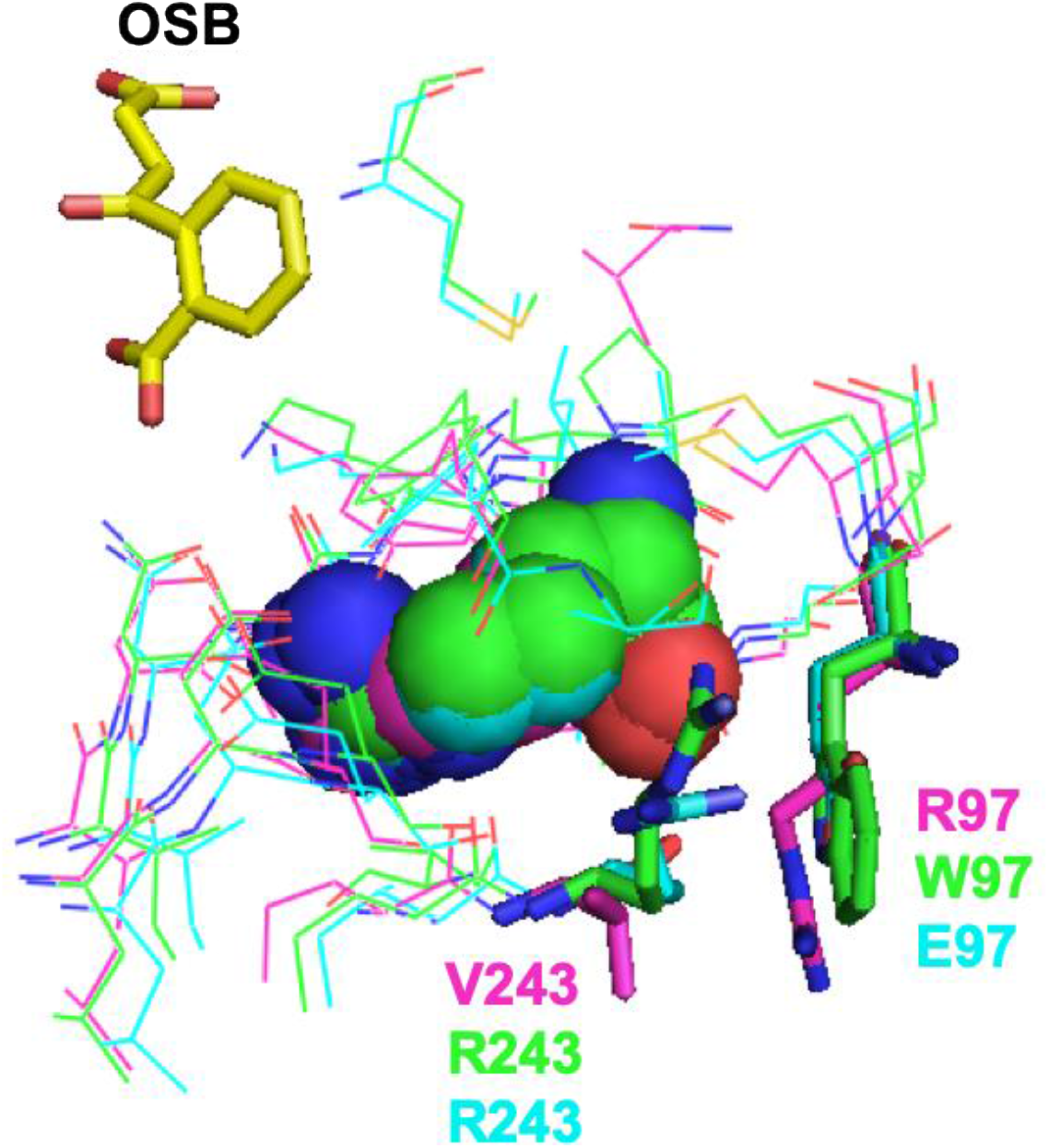
The local environments (within 5 Å) surrounding the R266 residue in EfNSAR/OSBS (PDB 1WUE, shown in green) [10], LiNSAR/OSBS (PDB 1WUF, shown in cyan) [10], and AmyNSAR/OSBS (PDB 1SJB, shown in magenta) [18]. R266 is shown in spheres. Residues 97 and 243 are labeled corresponding to the colors of the structures. OSB from 1SJB is shown in yellow.

### The roles of R266 in the Dipeptide Epimerase family of the MLE subgroup

The enolase superfamily includes several other protein families with racemization activity. For example, the muconate lactonizing enzyme (MLE) subgroup of the enolase superfamily contains the OSBS family, dipeptide epimerase (DE) family, 4R-Hydroxyproline Betaine 2-Epimerase family, and MLE 1 and 2 families (Figure 4). The members of the MLE subgroup contain the conserved catalytic lysine residues on the end of the second and sixth β-strands [19].

**Figure 4.**
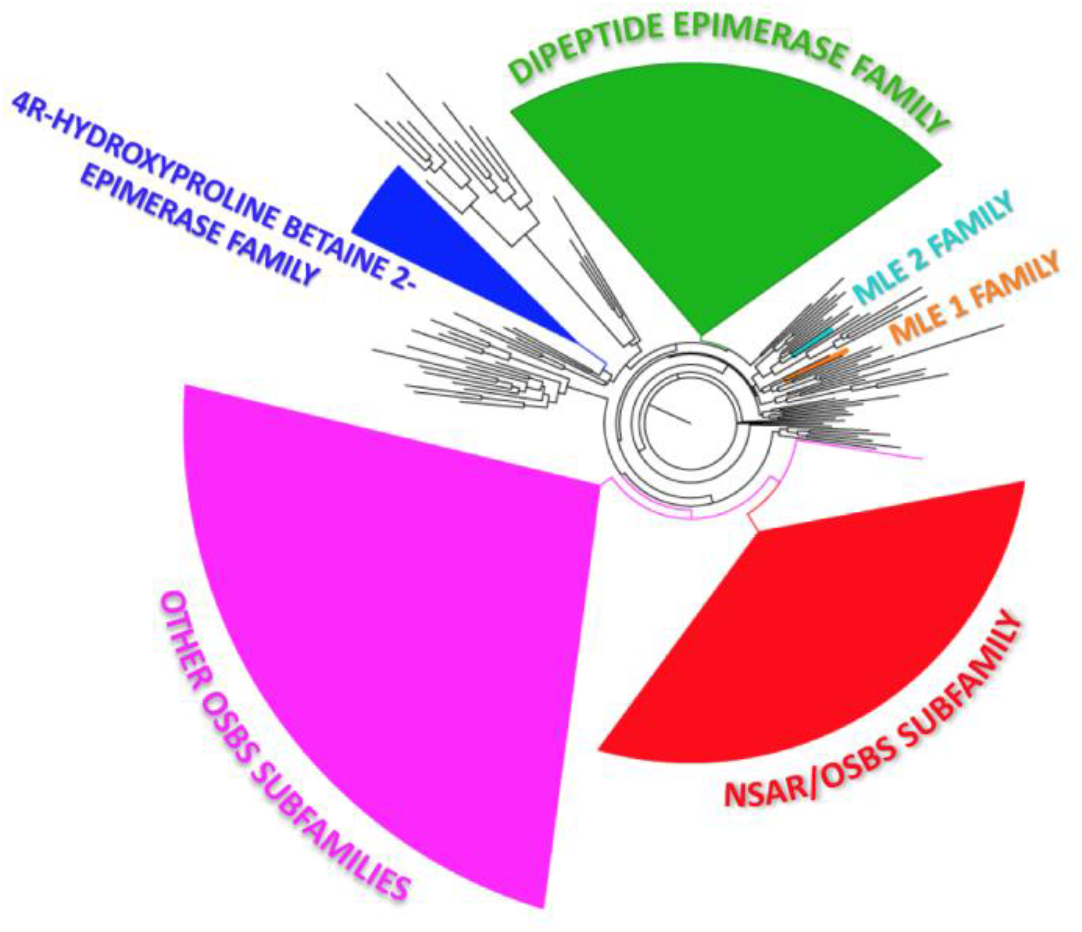
Phylogenetic tree of the MLE subgroup of the enolase superfamily. The MLE subgroup of the enolase superfamily contains the OSBS subfamilies (shown in magenta), the NSAR/OSBS subfamily (shown in red), the dipeptide epimerase (DE) family (shown in green), the 4R-Hydroxyproline Betaine 2-Epimerase family (shown in blue), and the muconate lactonizing enzyme (MLE) 1 and 2 families (shown in orange and cyan, respectively).

The DE family follows a two-base, 1-1 proton transfer mechanism similar to the NSAR reaction (Figure 5). Sequence logo analysis indicates that enzymes from this family also have a positively charged residue at position 266, but it is a lysine instead of an arginine (Figure 6). The structure of *Bacillus subtilis* DE (BsDE) with the bound Ala-Glu substrate shows that the homologous K266 forms a hydrogen bond with the homologous metal binding residue D239 (PDBID: 1TKK) (Figure 5) [20]. Conservation of a positively charged residue at position 266 in the NSAR/OSBS subfamily and the dipeptide epimerase family suggests that R/K266 is important for 2-base, 1-1 proton transfer reaction in the members of the MLE subgroup.

**Figure 5.**
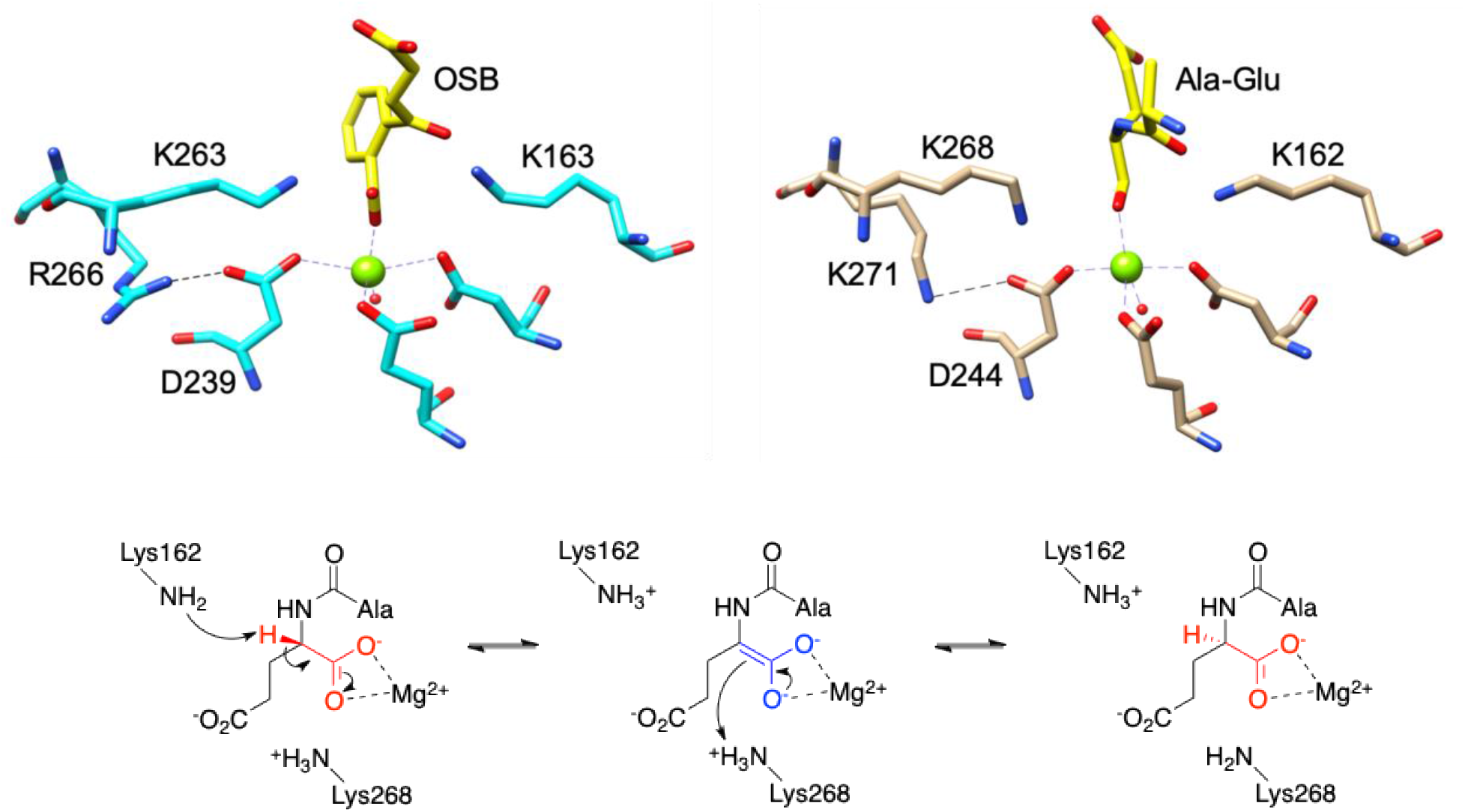
Comparison of the active sites of (A) AmyNSAR/OSBS (PDB 1SJB, [18]) and (B) BsDE (PDB 1TKK, [20]). (B) The mechanism of L-Ala-L/D-Glu dipeptide epimerase [21]. The divalent metal ion-stabilized enolate intermediate is shown in blue. The atoms that are rearranged during catalysis are shown in red. The numbering of the catalytic lysines are based on the sequence of *Bacillus subtilis* DE.

**Figure 6.**
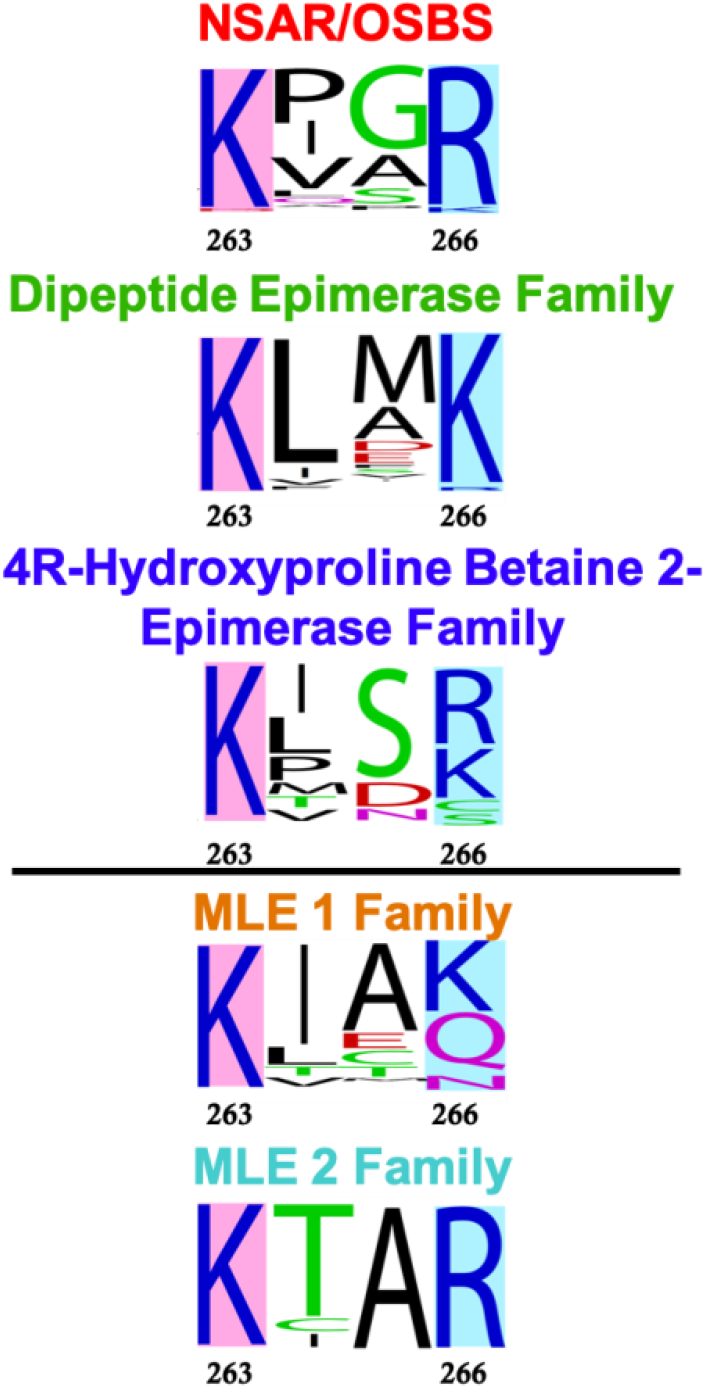
Sequence logos showing the conservation at positions 266 in some characterized families of the MLE subgroup, including the dipeptide epimerase family, the 4R-hydroxyproline betaine 2-epimerase family, and the MLE 1 and 2 families [22]. The letter size is proportional to the frequency at which each amino acid is found in the sequence alignment. The sequence numbering is relative to the *Amycolatopsis* NSAR/OSBS protein. Position 266 is highlighted in cyan. The catalytic K263 is highlighted in pink.

To test the hypothesis that the positively charged residue at the position equivalent to 266 is universally important for the 2-base, 1-1 proton transfer reaction, we also made the K250Q mutation in *E. coli* DE (EcDE) and K271Q mutation in *B. subtilis* DE (BsDE), which are homologous to R266 in AmyNSAR/OSBS. Like R266 in AmyNSAR/OSBS, K250 in EcDE and K271 in BsDE are second-shell amino acids and are not in contact with the substrate, suggesting they will not affect the binding of the substrate.

The K to Q mutation decreases *k*_cat_ of EcDE and BsDE by 44-fold and 93-fold, respectively, with no change in the K_M_ value in either enzyme (Table 5). This is consistent with the hypothesis that the positively charged residue at this position has the same effect in the NSAR and dipeptide epimerase reactions. We also measured the *k_ex_* values between the catalytic lysines and the alpha proton of the substrates to determine if this mutation impacts epimerase activity by affecting the reactivity of one lysine but not the other, as observed in AmyNSAR/OSBS (Table 6).

**Table 5:** Kinetic data for DE variants*

| **Table 5:** Kinetic data for DE variants* |  |  |  |
| --- | --- | --- | --- |
| | $K_M$ ($\mu\text{M}$) | $k_{\text{cat}}$ ($\text{s}^{-1}$) | $k_{\text{cat}}/K_M$ ($\text{M}^{-1}\text{s}^{-1}$) |
| **EcDE WT** | $340 \pm 60$ | $11.4 \pm 0.7$ | $(3.37 \pm 0.6) \times 10^4$ |
| **EcDE K250Q** | $385 \pm 85$ | $0.26 \pm 0.02$ | $(6.72 \pm 1.6) \times 10^2$ |
| **BsDE WT** | $452 \pm 120$ | $21.4 \pm 2$ | $(4.73 \pm 1.3) \times 10^4$ |
| **BsDE K271Q** | $508 \pm 165$ | $0.23 \pm 0.03$ | $(4.53 \pm 1.6) \times 10^2$ |
| *L-Ala-L-Glu was used as the substrate |  |  |  |

**Table 6:** Deuterium-Hydrogen exchange rate (*k_ex_*) of DE variants

| **Table 6:** Deuterium-Hydrogen exchange rate ($k_{ex}$) of DE variants | | | |
| --- | --- | --- | --- |
|  | **EcDE WT** | **EcDE K250Q** | **Fold change** |
| Ala-D-Glu and K151 | 70.4 s<sup>-1</sup> | 67.5 s<sup>-1</sup> | 1 |
| Ala-L-Glu and K247 | 34.5 s<sup>-1</sup> | 15.2 s<sup>-1</sup> | 2 |
|  | **BsDE WT** | **BsDE K271Q** | **Fold change** |
| Ala-D-Glu and K162 | 25.8 s<sup>-1</sup> | 0.78 s<sup>-1</sup> | 33 |
| Ala-L-Glu and K268 | 27.6 s<sup>-1</sup> | 0.13 s<sup>-1</sup> | 212 |

In BsDE, the K to Q mutation decreases the *k_ex_* values of the lysine of the 6^th^ beta strand, K268, more than those of the lysine on the 2^nd^ beta strand, K162, as expected. The reactivity of K162 decreases only by 33-fold while the reactivity of K268 decreases by 212-fold, approximately the same order of magnitude with the decrease of *k_cat_* by the K271Q mutation. However, in EcDE, the mutation only decreases the reactivity of K247 only by 2-fold while has no effect on the reactivity of K151. One possible explanation for the observed *k_ex_* values is that the *E. coli* strain used for DE protein expression also contains the native dipeptide epimerase, which could have co-purified with the mutant during purification. As a result, this experiment will be repeated by expressing and purifying the mutant in a DE knockout strain to ensure no contamination by native enzyme.

## Discussion

In this study, we examined the role of the second-shell amino acid R266 in the evolution of NSAR activity by examining the effects of the single substitution R266Q in other members of the NSAR/OSBS subfamily. We found that while the R266Q mutation decreases NSAR activity more than OSBS activity, as expected, in most NSAR/OSBS members, the differential effects of the R266Q substitution on NSAR and OSBS activities depend on differences in sequence and structural contexts among the enzymes. Our study on the second-shell amino acid R266 agreed with the idea that non-active site residues are more prone to epistasis than the catalytic residues [2]. In the NSAR/OSBS and DE families, mutation of the second-shell amino acid R266 has multiple phenotypic roles. We determined that R266 has an important role in catalysis in several enzymes, but in other cases it also has roles on protein stability and structure. For example, the effect of R266Q is quite specific in AmyNSAR/OSBS, and the phenotypic effects almost solely came from decreasing the reactivity of the catalytic K263. In other proteins, like LvNSAR/OSBS the mutation was very destabilizing and affected both activities as a result. These observations are in line with other studies that found that the same mutation will not have the same effect on stability of homologous proteins [3, 23]. We speculate that R266 has the same catalytic role in all of these enzymes, but deleterious effects on stability or overall structure caused by the R266Q mutation mask its underlying function to modulate the reactivity of K263. It is possible that saturation mutagenesis at this position could uncover a different mutation that has less effect on protein stability, revealing the more specific phenotypic effect of R266 on NSAR reaction specificity.

We demonstrated that both NSAR/OSBS and DE enzymes require a positively charged residue at the position equivalent to R266 in AmyNSAR/OSBS to carry out the 2-base, 1-1 proton transfer racemase/epimerase reactions, because mutations to glutamine this site significantly decreased their activities, primarily by decreasing *k_cat_*. In several cases, measuring *k_ex_* illustrated that decreased activity was due to a deleterious effect on the reactivity of K263. This observation supports the idea that gaining R/K266 was a pre-adaptive feature which enabled the emergence and evolution of racemase/epimerase activity. However, gaining R/K266 does not guarantee the enzymes will carry out racemase/epimerase activity. Interestingly, most enzymes from the MLE 1 and 2 families also have a positively charged residue at position 266, although they are not known to have racemase or epimerase activity (Figure 6). In MLE, the lysine on the 2^nd^ β-strand acts as an acid catalyst to protonate the enolate intermediate to yield mucolactonate. On the other hand, the lysine on the 6^th^ β-strand is closer to the carboxylic moiety of the product and presumably assists the stabilization of the enolate intermediate [24]. The role of this lysine on the 6^th^ β-strand is similar to that in the enzymes of the non-promiscuous OSBS subfamilies, which do not have a positively charged residue at position 266.

Examining the structure of mandelate racemase (MR), another member of the enolase superfamily, showed that MR enzymes also have a positively charged residue at the position homologous to 266, which is K273 (Figure 7). MR reversibly catalyzes the racemization of *R*- to *S*-mandelate, in a mechanism similar to NSAR and DE. In MR, while the conserved K166 on the end of the β2 strand is one catalytic acid/base, and H297 in the H297-D270 dyad is the other catalytic base. D270 has been shown to be important to assist H297 in catalysis [25]. The homologous lysine is in close proximity to D270 and a little further away from H297, but it could remotely interact with and potentially assist H297 in catalysis. The role of this lysine in MR activity has not been explored, and more evidence, including experimental characterization and mechanistic determination are necessary to test the role of this lysine in MR. However, a recent study showed that Y137, which is remote from H297, is essential for catalysis because it can modulate the pKa of both K166 and H297 by influencing the electrostatic environment of the whole active site [26].

**Figure 7:**
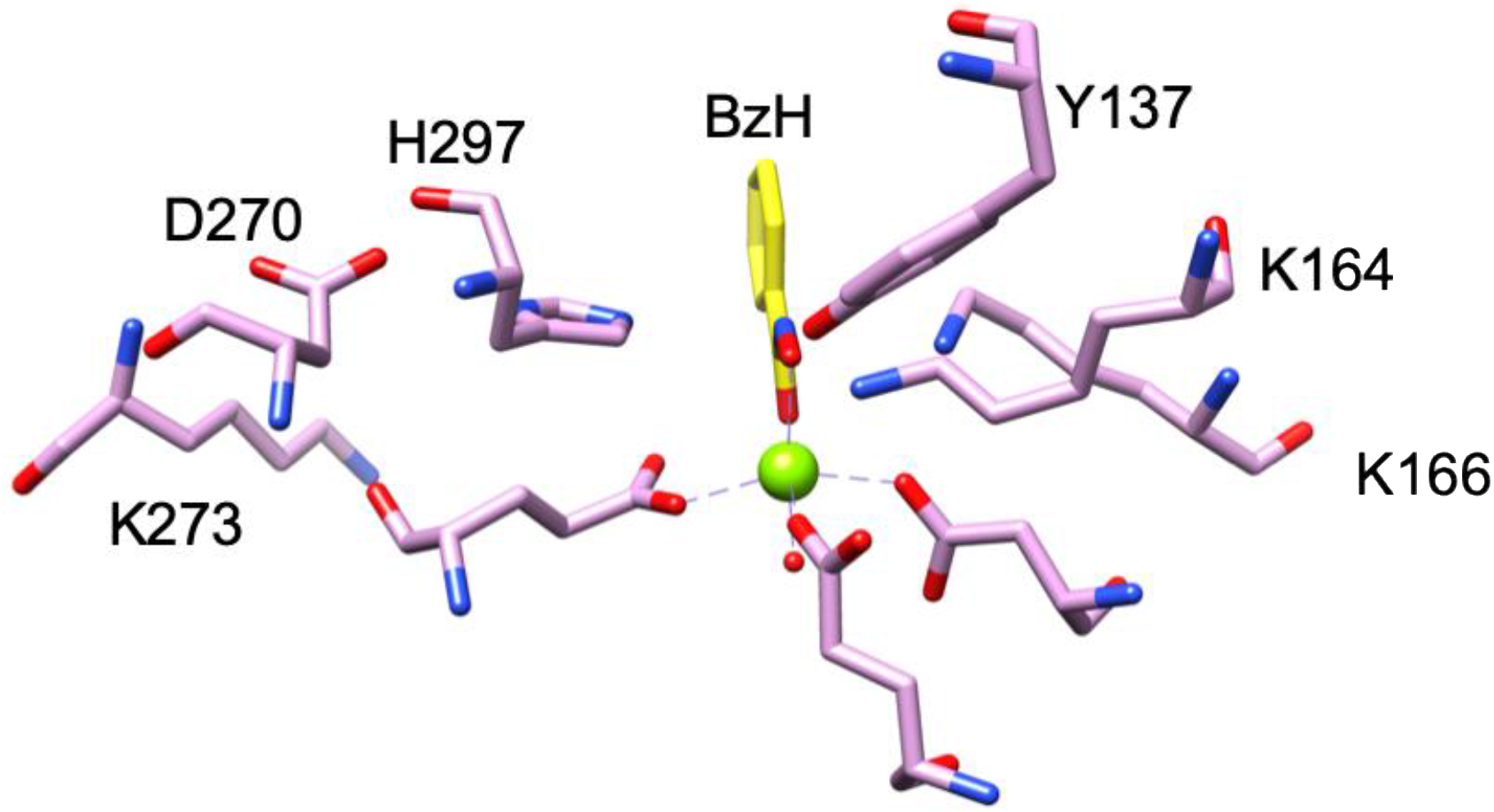
The active site of mandelate racemase (PDB ID 3UXK, [27]). The catalytic dyad H297-D270 is in close proximity to K273, which is homologous to R266 in AmyNSAR/OSBS. The catalytic triad Y137-K164-K166 is also shown. The intermediate/transition state analog benzohydroxamate (BzH) is shown in yellow.

An intriguing question is that whether an arginine or a lysine at this position can be used as a predictive tool for racemase/epimerase activity. Functional prediction and assignment are important yet difficult tasks for uncharacterized enzyme families. Many sequences with similar structures have similar functions, but some proteins with very similar structures have completely different functions [28]. On the other hand, enzymes with the same functions can have very different structures [29]. Functional prediction and annotation of enzyme superfamilies still remain a challenge even though there are powerful and advanced tools involving genomics, proteomics and metabolomics [30]. We showed that a positively charged amino acid at the position equivalent to 266 is important for racemase/epimerase activity in two members of the MLE subgroup of the enolase superfamily. Furthermore, the mandelate racemase enzymes from the enolase superfamily also have a lysine at this position, although its function is unknown. All characterized families with racemase/epimerase activity have a positively charged residue at position 266. However, not all the members of the MLE subgroup (and the enolase superfamily) have been experimentally characterized. With the data presented here, we can potentially use a positively charged amino acid at the position equivalent to 266 as an indicative tool for functional prediction and assignment in unassigned members of the enolase superfamily. We can compare the sequence alignment of the MLE subgroup to identify uncharacterized proteins with a positively charged amino acid at the position equivalent to 266. A positively charged residue at position 266 could suggest that racemase/epimerase activity is likely to be the biological function of the enzymes, providing a foothold for experimental characterization and mechanistic determination.

AUTHOR INFORMATION

## Funding

This work was funded by National Institutes of Health Award R01-GM124409, National Science Foundation CAREER Award 1253975, and the Welch Foundation Grant No. A-1991-20190330 (Principal Investigator, Dr. Margaret E. Glasner).

## Notes

## ACKNOWLEDGMENTS

We thank members of D.P.T.’s Ph.D. advisory committee for helpful comments.

## ABBREVIATIONS

OSBS: *o*-succinylbenzoate synthase
NSAR: *N*-succinylamino acid racemase
EfNSAR/OSBS: *Enterococcus faecalis* NSAR/OSBS
RcNSAR/OSBS: *Roseiflexus castenholzii* NSAR/OSBS
LvNSAR/OSBS: *Lysinibacillus varians* NSAR/OSBS
LiNSAR/OSBS: *Listeria innocua* NSAR/OSBS
*k_ex_*: exchange rate
DE: Dipeptide Epimerase
NSPG: *N*-succinylphenylglycine
SHCHC: 2-succinyl-6-hydroxy-2,4-cyclohexadiene-1-carboxylate
DSF: Differential Scanning Fluorimetry

## Notes

### Competing Interest Statement

The authors have declared no competing interest.

